# A Generic Numbering Scheme for TMEM16 Scramblases

**DOI:** 10.64898/2026.09.01.748741

**Authors:** Shuai Yan, Harel Weinstein

## Abstract

The TMEM16 family of calcium-activated phospholipid scramblases (CaPLSs) and chloride channels (CaCCs) performs diverse physiological functions that include regulation of blood coagulation and apoptotic signaling, through a shared ten-transmembrane-helix (TM) architecture organized around a hydrophilic lipid-translocating groove. Mechanistic studies of TMEM16 family members have been hampered by the absence of a unified positional reference framework that would permit direct comparison of structurally equivalent residues across paralogs with different sequence numbering systems. Here we introduce a generic numbering scheme for TMEM16 scramblases (GNS-TMEM16), modeled on the Ballesteros & Weinstein system established for class A G protein-coupled receptors. A reference alignment (TMEM16-RA) was constructed from twelve human and mouse TMEM16 scramblases (TMEM16C/D/E/F/G/J) using structure-based ClustalW alignment of the ten TM helices. From this alignment, a TM-specific reference residue (TsRR) was identified for each helix by hierarchical application of three criteria: (1) 100% conservation in the core TMEM16-RA; (2) conservation in an augmented reference alignment (TMEM16-ARA) incorporating a group of phylogenetically more distant homologs composed of nhTMEM16, afTMEM16, TMEM16K, TMEM16A, and TMEM16B; and (3) structural and functional considerations, including helix-perturbing character, groove localization, conserved motif membership, and central TM position. The resulting ten TsRRs are Y1.50, W2.50, R3.50, E4.50, F5.50, P6.50, E7.50, D8.50, W9.50, and E10.50, and are illustrated in mTMEM16F. Each residue is assigned the identifier **N.m(k)**, where **N** is the TM number, **m** is the position relative to the TsRR (for which m = 50), and **k** is the absolute sequence number. Loop residues receive dual identifiers referenced to the TsRRs of both flanking helices. Application of the GNS-TMEM16 is illustrated with the comparisons of the groove-opening measurements using pairwise distances between residues identified by their **N.m** indices to be corresponding across mTMEM16F, afTMEM16, and nhTMEM16. The results bring to light the advantages of corresponding residues identification in different TMEM16 proteins and show that the mammalian scramblase undergoes substantially larger separation at the extracellular groove entrance than either fungal homolog. Comparison of mutagenesis data guided by N.m correspondence shows at the conserved (E3.55,R6.26) salt-bridge locus, Ala substitution reduces activity more than 100-fold in nhTMEM16 but less than 2-fold in afTMEM16, illustrating that the GNS identifies structural equivalence of position without implying functional equivalence of the residue, which is a distinct advantage of GNS in providing mechanistic interpretation across paralogs. Also described is a protocol for extending the GNS-TMEM16 to uncharacterized protein sequences, including AlphaFold-predicted models, using structural superposition to mTMEM16F. Thus, the presented GNS-TMEM16 provides a stable positional reference for the integration and comparative analysis of structural, computational, and functional data across the TMEM16 family, utilizing a construction strategy applicable to yet other polytopic membrane protein families sharing a common transmembrane fold.

## Introduction

The sustained interest in the TMEM16 family of proteins reflects the diversity of the physiological functions they perform while sharing closely similar structures. Thus, the characterization of the mammalian TMEM16 family members (TMEM16A, TMEM16F, TMEM16K) and of members of the fungal TMEM16 family (afTMEM16 and nhTMEM16) (Lee et al., 2016; Malvezzi et al., 2013) has determined their roles as calcium-activated phospholipid scramblases (CaPLSs) (Duran & Hartzell, 2011, p. 2009; Pifferi et al., 2009; Schroeder et al., 2008; Yang et al., 2012) and/or calcium-activated chloride channels (CaCCs) (Bushell et al., 2019). The physiological functions of CaPLSs include modulation of the composition and configuration of the cell membrane. For example, TMEM16 CaPLSs increase the exposure of phosphatidylserine (PS) in the extracellular leaflet—a feature associated with physiological processes including blood coagulation and the removal of apoptotic cells (S. C. Le et al., 2021). In many cases, the CaCC and CaPLS functions were shown to involve similar structural motifs of the TMEM16 proteins, yet systematic comparisons across family members have not been performed. Comparisons of structure–function relationships are essential for discerning the distinct functional mechanisms underlying the diverse physiological roles of these proteins. To facilitate such comparisons, we have developed a generic numbering scheme (GNS) for members of the TMEM16 family of CaPLSs and CaCCs, which also enables the inclusion of other protein families with similar folding architectures in such analyses.

The significant advantages offered by a GNS for elucidating the functional mechanisms of membrane proteins are amply demonstrated by the widespread adoption of the well-established GNS developed for class A G protein-coupled receptors (GPCRs) (Ballesteros & Weinstein, 1995). An extensive literature illustrates how this GNS enables the study of functional mechanisms across different GPCRs within a unified framework, by allowing direct recognition and functional comparison of corresponding structural motifs in different proteins despite differences in their sequence numbering and composition (Bhattacharya et al., 2016; Billesbølle et al., 2023; Conflitti et al., 2025; White et al., 2018; Zhang et al., 2025; Zhou et al., 2019). Results from investigations of the structure–function properties of TMEM16 family members have revealed features that suggest the feasibility of an analogous treatment. Specifically, TMEM16 CaPLS and CaCC family members share a relatively high degree of sequence homology and a common fold architecture, and detailed studies have shown that these similarities extend to structural elements and their dynamic properties underlying their different structural mechanisms (Picollo et al., 2015).

TMEM16 CaPLS and CaCC proteins function as homodimers (**Figure 1)** activated by the binding of two or three Ca2+ ions (Brunner et al., 2014; S. C. Le & Yang, 2020; Peters et al., 2018). Their main structural landmarks include a central cavity formed by the two protomers and a subunit cavity— commonly referred to as the groove—formed by transmembrane helices (TM) 3, TM4, and TM6 (Brunner et al., 2014) in each protomer (**Figure 1**). In the open-state conformation capable of translocating phospholipids or ions, TM4 and TM6 separate within each protomer, which further exposes to the membrane a hydrophilic groove lining composed of polar and charged side chains.

**Figure 1.**
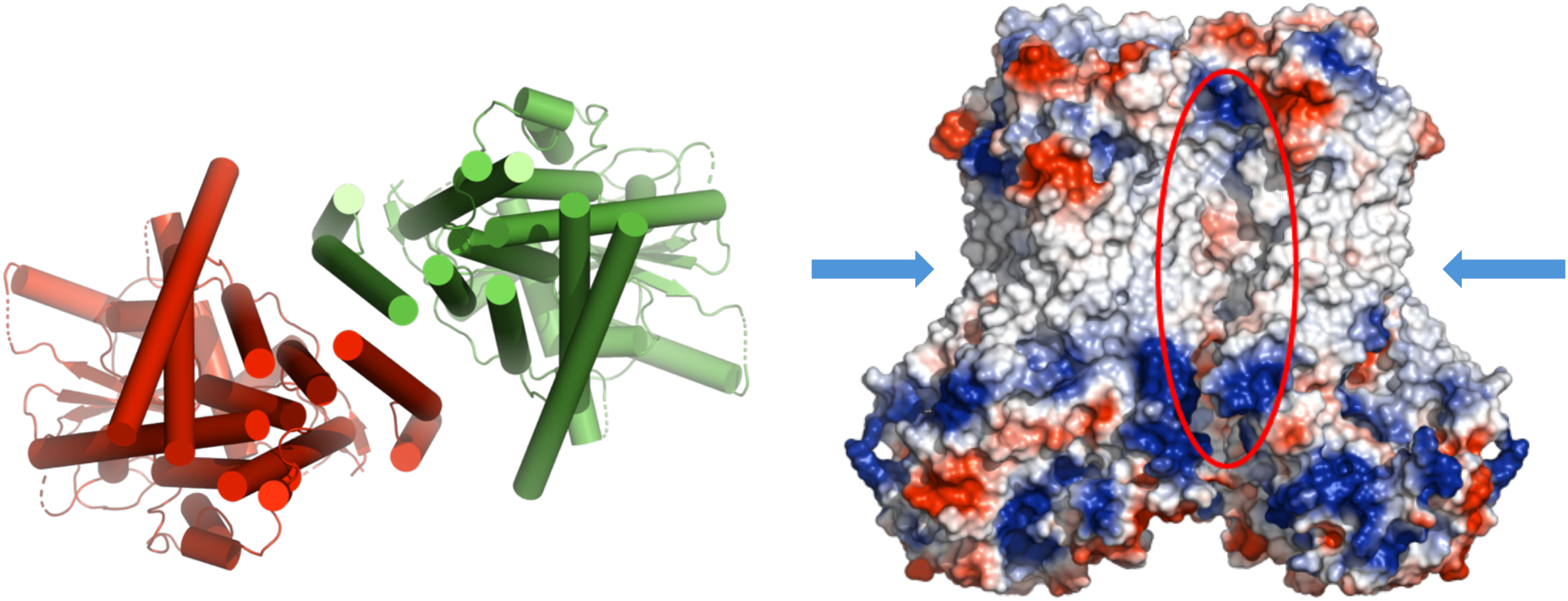
Structural representation of key features of the mTMEM16F dimer. **(Left Panel)** Representation of the 3D architecture of the helical domains of the homodimer. The two protomers are rendered in different colors (in red and green) with TMs in cartoon representation. The view is from the extracellular side, perpendicular to the membrane (omitted for simplicity). **(Right Panel)** Volume rendering of the homodimer with surface colored to represent polar regions (red and blue) and non-polar regions (in white). The dimer cavity formed by TM3 and TM10 is indicated by the red ellipse, and the subunit cavities (grooves) facing the surrounding membrane are indicated by light blue arrows for each protomer.

Evidence from structural studies and functional assays shows that the open-groove conformation is favored upon Ca2+ binding at the conserved sites, which potentiates the translocation function. In many, but not all, cases, this Ca2+-dependent activation promotes groove opening (Alvadia et al., 2019; Falzone et al., 2022). Despite this commonality, sequence and structural differences among TMEM16 family members in other regions of the protein are associated with additional functional distinctions, including differences in cellular localization and lipid type preference, that extend beyond the established CaCC/CaPLS classification (Suzuki et al., 2013).

The molecular mechanisms of the CaPLSs have been studied extensively using a small number of fungal TMEM16 homologs, predominantly nhTMEM16 and afTMEM16. As mechanistic differences were identified among the fungal scramblases, and as comparisons were extended to mammalian scramblases such as mTMEM16F, it became increasingly difficult to interpret sequence differences between paralogs in terms of the underlying structural distinctions. As demonstrated for the GPCR family, the ability to compare functional properties across homologous membrane proteins, and to infer mechanistic generalizations in a structural context can be greatly enhanced by a GNS because it enables direct recognition of structurally equivalent positions within different sequence numbering systems (Ballesteros & Weinstein, 1995).

Here we present such a numbering scheme for TMEM16 scramblase residues, analogous to the established GNS for class A GPCRs (Ballesteros & Weinstein, 1995). We show that starting from structure-based sequence alignments of the transmembrane helices of highly homologous TMEM16 proteins—in which TMs are structurally superposed to prevent spurious insertions or deletions within helical segments—it is possible to construct a reference alignment (TMEM16-RA) that parses the corresponding structural elements (transmembrane helical segments and TM-connecting loops) and identifies the 100%-conserved residues within each TM. As in the GPCR system, one such highly conserved residue per TM is assigned the locus number 50, providing a TM-specific anchor from which all other residues in that helix are numbered relative to that reference position (denoted N.50, where N is the TM number). Residues on the N-terminal side of the reference are numbered N.49, N.48, etc., while those on the C-terminal side are N.51, N.52, etc. As part of presenting and illustrating the construction of the GNS-TMEM16, we also describe a protocol for incorporating newly encountered TMEM16 sequences into the numbering scheme.

## Results

### 2.1 Construction of the TMEM16 Numbering Scheme

#### 2.1.1 Sequence similarity and selection of sequences for the reference alignment

Construction of the TMEM16 numbering scheme exploits the high degree of sequence and structural similarity among experimentally characterized TMEM16 scramblases. The phylogenetic tree in **Figure 2** indicates the close evolutionary relationships among human and mouse TMEM16 proteins C, D, E, F, G, and J, all of which are considered to function as phospholipid scramblases (Di Zanni et al., 2018; Petkovic et al., 2020; Suzuki et al., 2013). The evolutionary history was inferred using the Maximum Likelihood method with the Le Gascuel 2008 substitution model (S. Q. Le & Gascuel, 2008).

**Figure 2.**
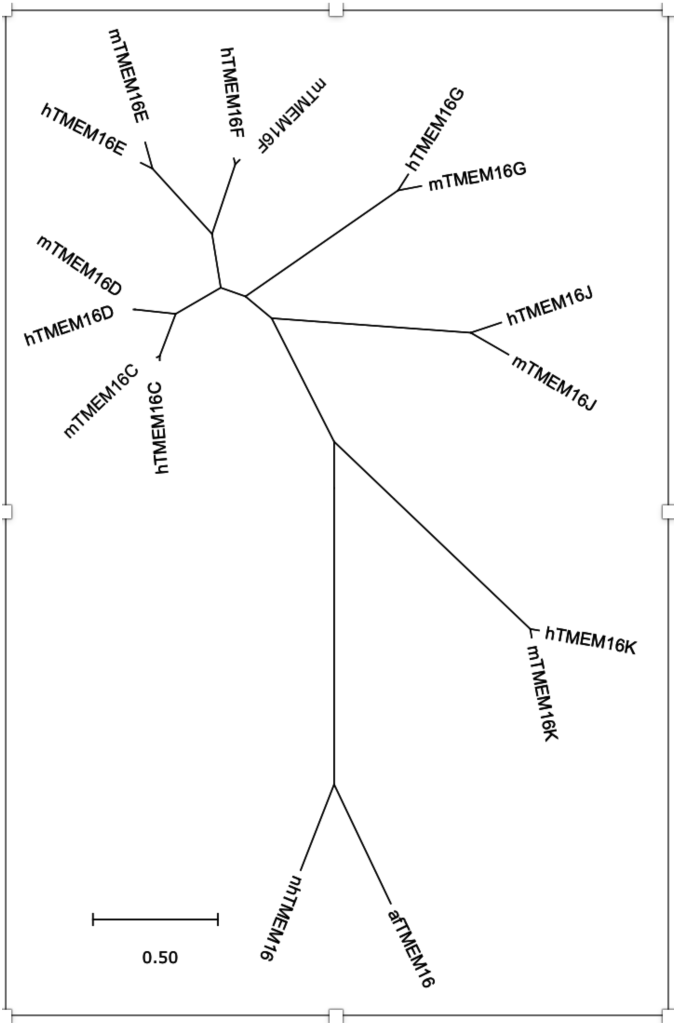
Phylogenetic tree of TMEM16 homologs. Evolutionary analyses were conducted in MEGA11 (Tamura et al., 2021) using the Maximum Likelihood method and Le Gascuel 2008 model (S. Q. Le & Gascuel, 2008). The tree with the highest log-likelihood (−11524.93) is shown to scale, with branch lengths in substitutions per site. Sixteen amino acid sequences were analyzed; sites with less than 95% coverage were excluded. The final dataset contained 541 positions.

Initial trees for the heuristic search in the construction of the tree were obtained by applying the Neighbor-Joining and BioNJ algorithms to a matrix of pairwise distances estimated with the JTT model (Jones et al., 1992), and the topology with the superior log-likelihood value was selected. Evolutionary rate differences among amino acid sites were modeled with a discrete Gamma distribution (5 categories; +G parameter = 4.6565). The default partial-deletion option in MEGA11 (Tamura et al., 2021) was applied, excluding sites with less than 95% coverage across all sequences. Sites containing gaps, missing data, or ambiguous characters were counted against coverage. The final dataset comprised 541 positions. Inspection of **Figure 2** shows that the lower part consists of 4 sequences that are distant from the larger group of TMEM16 mammalian scramblases clustered in the upper part of the phylogenetic tree. This group of distal sequences is composed of the fungal homologs afTMEM16 and nhTMEM16 and includes as well the TMEM16K expressed in humans and mice. We reasoned that construction of an initial reference alignment of TMEM16 proteins (TMEM16-RA) including only the residues in the upper cluster, would yield a stricter conservation criterion for the numbering scheme. It may limit candidate positions for the “TM-specific reference residues” (TsRR) in most TMs to 100% conserved ones. Therefore, we selected for the initial TMEM16-A the homologs with best structural characterization among those included in the upper part of the phylogenetic tree in Figure 2. This includes the sequences of human and mouse TMEM16 proteins C, D, E, F, G, and J (Di Zanni et al., 2018; Petkovic et al., 2020; Suzuki et al., 2013).

The sequence alignment was performed in MEGA11 (Tamura et al., 2021; Thompson et al., 1994) following the protocol described in (Hall, 2013) (see alignment protocol and parameters in the Methods Section). The full TMEM16-RA alignment is shown in **Supplement Figure 1.** Inspection of this **Suppl.Fig. 1** reveals that TMs 6, 7, and 8 — which participate in binding two Ca^2+^ ions (Brunner et al., 2016) — are the most conserved among the aligned scramblases. TM3, which lines the protomer groove, is among the least conserved TMs.

### 2.2 Criteria for Selection of TM-Specific Reference Residues (TsRRs)

#### Criterion 1: Selection of a TsRR from the TMEM16-RA

For a TM-specific reference residue (TsRR) to be applicable across all TMEM16 scramblases, it must be present in most or all homologs. This requirement points to 100% conservation within the reference sequence alignment as the primary criterion for TsRR selection. Inspection of the TMEM16-RA in **Suppl. Fig. 1** confirms that every TM contains at least one residue that meets this criterion.

Because of the high overall sequence similarity among the aligned homologs, application of Criterion 1 alone produced multiple TsRR candidates per TM. Two additional stringency criteria were therefore introduced.

#### Criterion 2: Selection of a TsRR from the augmented reference alignment (ARA)

To reduce the number of candidates identified by Criterion 1, we augmented the core TMEM16-RA with several of the phylogenetically more distant TMEM16 homologs. These include the mammalian TMEM16K and the fungal scramblases afTMEM16 and nhTMEM16, for all of which experimental structures are available, as well as the well-characterized calcium-activated channel homologs TMEM16A and TMEM16B. This augmented reference alignment (ARA) revealed subtype-specific conserved positions, as well as positions shared between scramblases and channels, thereby increasing the discriminating power of the selection procedure.

Application of Criterion 2 proceeds as follows. The set of candidate loci identified in in the initial TMEM16-RA for each TM by Criterion 1, is reduced by selecting the position that is also conserved in at least one homolog in the ARA (i.e., a “consensus conservation”). This conservation is sought in a specific order: first among the additional mammalian scramblases, then among the fungal scramblases, and finally among the CaCC sequences in the ARA. The first consensus conservation identified in this order is used to narrow the candidate set.

We found that despite applying both Criteria 1 and 2, TMs 1–3, 5–8, and 10 still presented more than one candidate for TsRR in several TMs, due to the high overall conservation of the protein family. Criterion 3 was therefore introduced to resolve the remaining ambiguities.

#### Criterion 3: Consideration of structure–function information

From among the candidates satisfying Criteria 1 and 2, the TsRR is selected as the residue with one or more of the following structural or functional attributes:

i. The residue is a putative helix de-constructor (e.g., Pro, Gly, Tyr, or Ser (Ballesteros et al., 2000; S.P. Sansom & Weinstein, 2000; Tooze, 1998)). Conservation of such residues within a TM implies preservation of local structural and functional properties.
ii. The residue is located in the groove region, indicating a likelihood of interaction with phospholipid substrates during scrambling.
iii. The residue is part of a highly conserved sequence motif.
iv. The residue is positioned near the center of the TM, which minimizes the impact of differential helix lengths on its relative position across homologs during structural alignment.

### 2.3 Hierarchical Application of Criteria 1–3: Illustrative Examples

The hierarchical application of the three criteria to the ARA is illustrated for two representative TMs— TM2 and TM8—for which several rounds of filtering were required (Figure 3).

**Figure 3.**
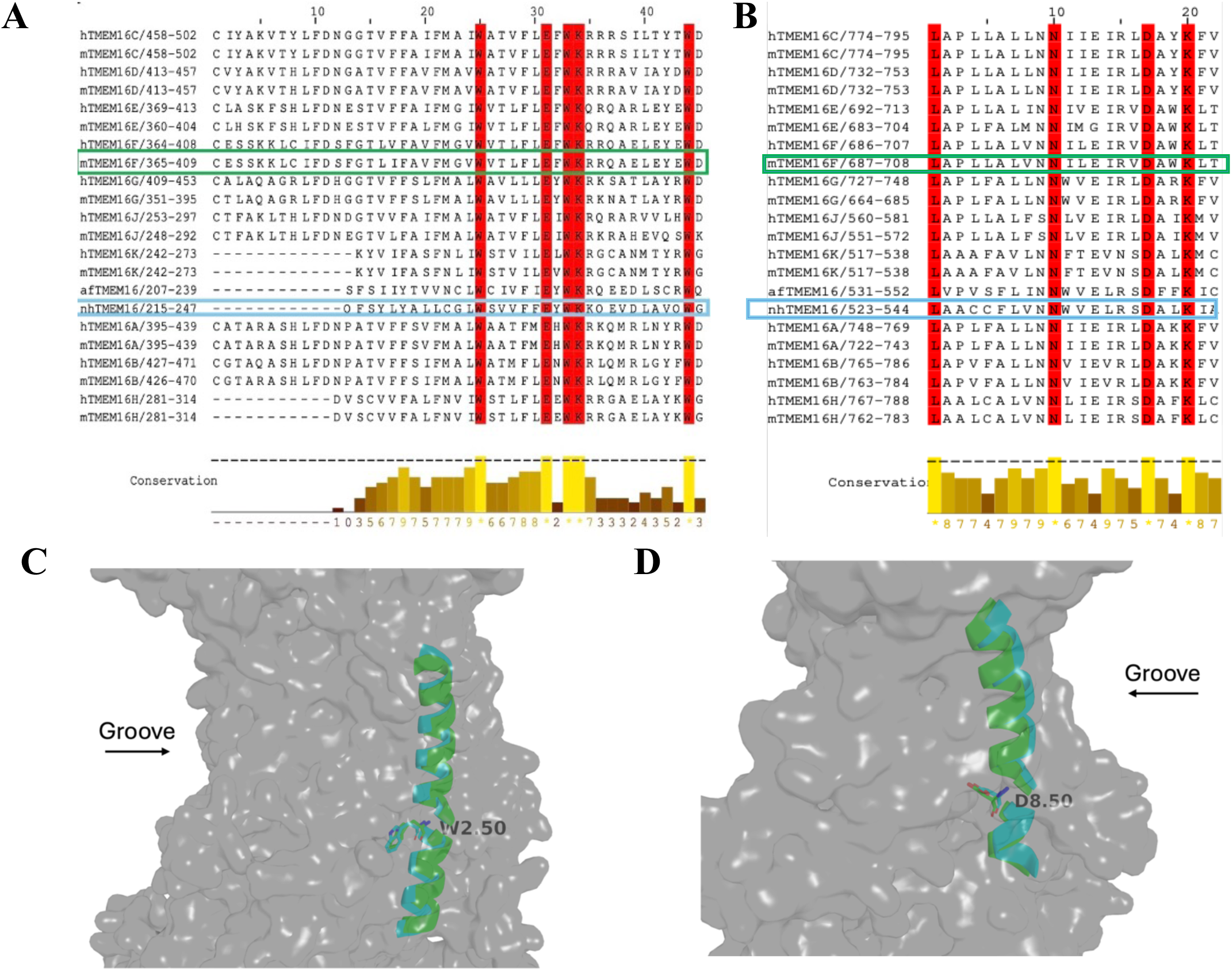
Steps in the application of Criteria 2 and 3 to identify the TsRRs for TM2 and TM8. Sequence alignments generated with Jalview (Waterhouse et al., 2009) of TM2 (**in A**), and of TM8 **(B)** in the ARA. Residues satisfying Criterion 1 (100% conservation) are highlighted *in red*. To select the TsRR among the multiple candidates in each TM, Criteria 2 and 3 are applied as described in the text. **Panels C and D** show the resulting selection of TsRRs for TM2 and TM8. The grey background in each panel is a 3D rendering of the RMSD the same superposition of the two TMEM16 structures: mTMEM16F (PDB: 6QPB) and nhTMEM16 (PDB: 6QM4) in their closed states, but with the view rotated to show separately the two TMs, highlighted in color. **Panel C** presents the superposition of TM2 in mTMEM16F (***green)*** and of nhTMEM16 (in ***cyan).* Panel D** highlights the superposition of TM8 of the two proteins in the same color scheme. The sequences of TM2 and TM8 in each of the two proteins are highlighted in the alignment Panels (**A** and **B)** by boxes in the corresponding colors of the TMs (green and cyan). **Note the positional coincidence of the corresponding 2.50 and 8.50 residues in the mammalian and fungal proteins.**

For TM2 (**Figure 3a**), Criterion 1 identified five candidate positions (in sequence order: Trp, Glu, Trp, Lys, Trp). None of these residues has been specifically implicated in a functional role, and none is located in the groove region; their spatial positions are also invariant between open- and closed-groove conformations in available cryo-EM structures. Therefore, Criterion 2, and elements 3(i) to 3(iii) in Criterion 3, provided no further discrimination. Only Criterion 3(iv) was informative for the refinement of the TsRR selection: the two terminal Trp residues in TM2 were eliminated as candidates because they lie far from the conserved central motif of TM2. Among the three remaining candidates (Glu, Trp, Lys), which are equivalent under all applicable sub-criteria, the one nearest the midpoint of TM2 is Trp, which was selected as the TsRR.

For TM8 (**Figure 3b)**, sequence and conservation considerations narrowed the candidates to four residues (in sequence order: Leu, Asn, Asp, Lys). To evaluate Criterion 3(ii), open- and closed-groove conformations of experimental structures were superposed in PyMOL (Schrödinger, LLC, 2015), and the solvent accessibility of each candidate was assessed relative to the groove surface. Atoms of candidate residues that protrude beyond the groove surface in the closed conformation within their van der Waals radii (Bondi, 1964) were scored as groove-lining. Both Asp and Lys satisfy this criterion.

Between these two, Asp is positioned closer to the center of TM8; selecting it as the TsRR therefore yields lower root-mean-square deviation (RMSD) values when TM8 segments from different TMEM16 subtypes are structurally aligned on the reference position. Accordingly, the Asp was chosen as the TsRR for TM8.

### 2.4 Description of identifiers in the Generic Numbering Scheme

The GNS for TMEM16 scramblases assigns to each residue in a transmembrane helix a three-part identifier of the form ***N.m(k)***, where: **N** is the TM segment number in the structure of the protein; **m** is the position of the residue relative to the TsRR of that TM, which is assigned the value 50; and **k** is the residue number in the amino acid sequence of the specific protein. For a specific protein, these three numbers are written in the form ***N.m(k)***, preceded by the one-letter code of the specific residue at that location. For example, in nhTMEM16 the residue glutamate E313 is located in TM3, five residues downstream (toward the C-terminus) from the TsRR of TM3, which is Thr308 (assigned the locus 3.50). Its full identifier is therefore *E3.55(313)* and the TsRR is *T3.50(308)*.

For residues in TM-connecting loops, the identifier encodes the distance from the TsRRs of both flanking TMs. For example, the identifier *D3.70/4.08(497)* for Asp497 in TMEM16F indicates that this residue lies in the loop between TM3 and TM4, 20 residues downstream from the TsRR of TM3 (position 3.50) and 42 residues upstream from the TsRR of TM4 (position 4.50); its sequence number is 497. Residues in non-TM segments of the N- and C-termini are excluded from the GNS.

Mutations are represented in the standard format: for example, substitution of Arg432 in nhTMEM16 with Trp is written as *R6.26(432)W*. This notation makes it straightforward to identify and compare mutations at structurally equivalent positions across different TMEM16 family members.

Table 1 lists the TsRR for each of the ten TMs, together with the corresponding residue identities and sequence positions in mTMEM16F. These reference residues are labeled in **Figure 4** on the closed-groove structure of mTMEM16F.

**Figure 4.**
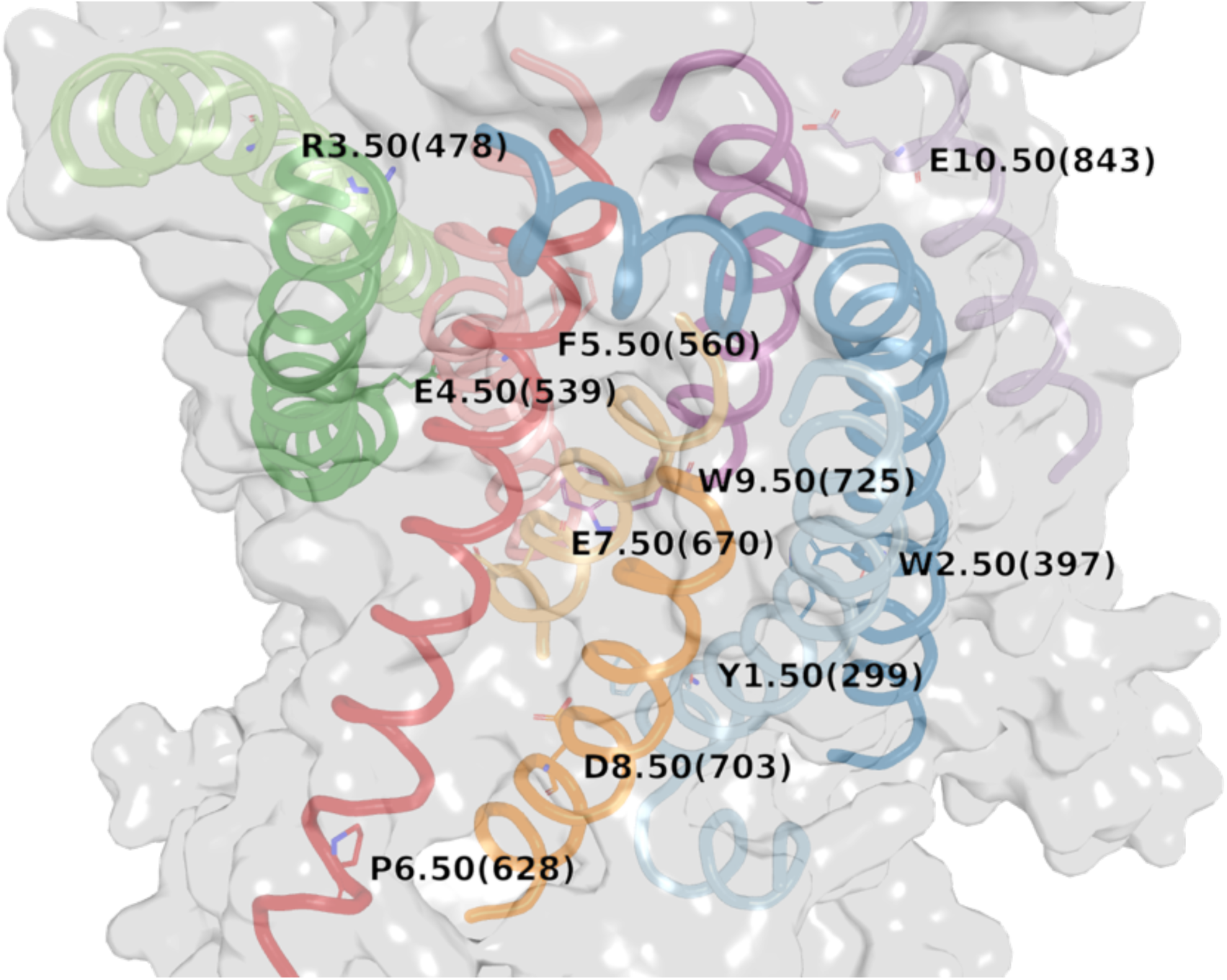
The GNS-TMEM16 numbering scheme illustrated on the structure of mTMEM16F (PDB: 6QPB). TM1–TM10 are shown with their respective TsRRs labeled. The groove is oriented to the left; the extracellular leaflet of the membrane (not shown) is at the top. TMs are colored distinctly: TM1, dark green; TM2, cyan; TM3, magenta; TM4, yellow; TM5, wheat; TM6, lemon yellow; TM7, slate blue; TM8, orange; TM9, light green; TM10, deep teal. In the reference residues, carbon atoms are colored to match their backbone ribbon, nitrogen atoms are blue, and oxygen atoms are red.

**Table 1.** TM-specific reference residues (TsRRs) identified from the TMEM16-RA and their corresponding positions in mTMEM16F.

| TM | Conserved residue | TsRR identifier | Residue no. in mTMEM16F | Full identifier in mTMEM16F |
| --- | --- | --- | --- | --- |
| 1 | Tyr | Y1.50 | 299 | Y1.50(299) |
| 2 | Trp | W2.50 | 397 | W2.50(397) |
| 3 | Arg | R3.50 | 478 | R3.50(478) |
| 4 | Glu | E4.50 | 539 | E4.50(539) |
| 5 | Phe | F5.50 | 560 | F5.50(560) |
| 6 | Pro | P6.50 | 628 | P6.50(628) |
| 7 | Glu | E7.50 | 670 | E7.50(670) |
| 8 | Asp | D8.50 | 703 | D8.50(703) |
| 9 | Trp | W9.50 | 725 | W9.50(725) |
| 10 | Glu | E10.50 | 843 | E10.50(843) |

### 2.5 Illustrations of GNS use applications

#### 2.5.1 Measuring Groove Opening

Khelashvili et al. (Khelashvili et al., 2022) selected pairs of groove-lining residues to characterize the groove-opening process in TMEM16F using molecular dynamics (MD) simulations. TMEM16F differs from the fungal TMEM16 scramblases in that it contains a helix-loop-helix cap in TM2, which influences the groove-opening mechanism. An open-groove conformation of TMEM16F had not yet been captured by cryo-EM, so Khelashvili et al. (Khelashvili et al., 2022) characterized the open-groove state from the analysis of the MD simulation trajectories with the time-lagged independent component analysis (tICA) method (Schultze & Grubmüller, 2021).

Applying the GNS to translate the residue pair identifiers used in mTMEM16F into equivalent positions in afTMEM16 and nhTMEM16, we measured the corresponding Cα–Cα distances between groove-opening pairs in the closed- and open-groove conformations of the two fungal scramblases. The nhTMEM16 measurements are shown as an example in **Figure 5**; results for all three proteins are compiled in Table 2.

**Figure 5.**
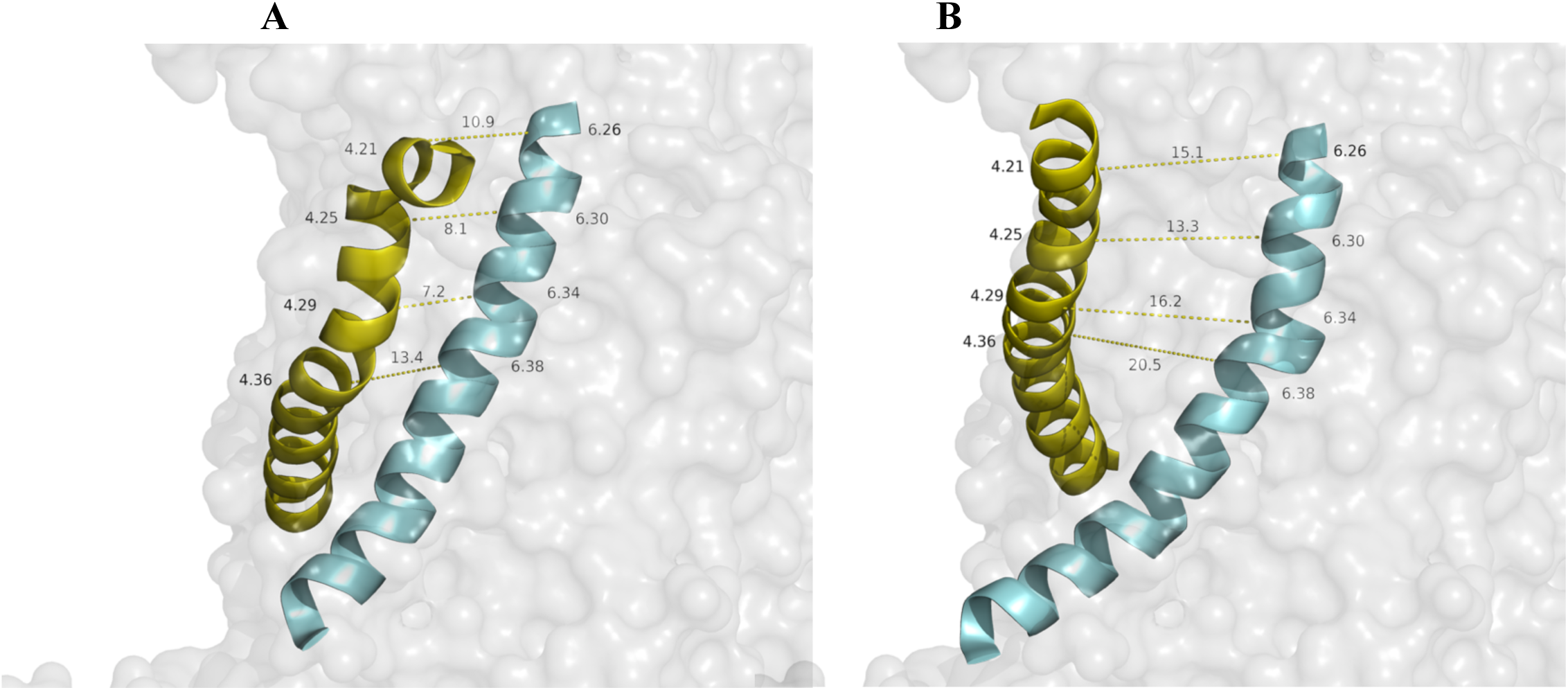
Cα–Cα distances between groove-lining residues in TM4 and TM6 of nhTMEM16. **(A)** Shows the closed-groove conformation, and **(B)** shows the open-groove conformation obtained from PDB entries 6QM4 (closed) and 4WIS (open), respectively; see also results for these states in (Khelashvili et al., 2019). TM4 is rendered in yellow, and TM6 in cyan. The data for key distances in Table 2 were obtained from the same measurements protocols applied to the structures of afTMEM16 using the PDB entries 6D7Z (closed) and 6EOH (open). The mTMEM16F data are from Figure S10 of Khelashvili et al. (Khelashvili et al., 2022).

**Table 2.** Groove-opening distances for TM4–TM6 residue pairs across three TMEM16 scramblases. mTMEM16F data are adapted from Figure S10 of Khelashvili et al. (Khelashvili et al., 2022). afTMEM16 was measured using PDB entries 6D7Z (closed) and 6EOH (open); nhTMEM16 was measured using PDB entries 6QM4 (closed) and 4WIS (open). Distances are Cα–Cα distances for the indicated residue pairs.

| Residue pair<br>(mTMEM16F) | GNS<br>identifier | mTMEM16F<br>($\Delta$=closed→open) | afTMEM16<br>($\Delta$=closed→open) | nhTMEM16<br>($\Delta$=closed→open) |
| --- | --- | --- | --- | --- |
| S510–E604 | 4.21–6.26 | ~6.5 Å → ~14.0 Å<br>($\Delta \approx 7.5$ Å) | 12.7 Å → 14.3 Å<br>($\Delta \approx 1.6$ Å) | 10.9 Å → 15.1 Å<br>($\Delta \approx 4.2$ Å) |
| S514–Q608 | 4.25–6.30 | ~5.0 Å → ~13.5 Å<br>($\Delta \approx 8.5$ Å) | 7.6 Å → 12.0 Å<br>($\Delta \approx 4.4$ Å) | 8.1 Å → 13.3 Å<br>($\Delta \approx 5.2$ Å) |
| F518–I612 | 4.29–6.34 | ~5.5 Å → ~14.0 Å<br>($\Delta \approx 8.5$ Å) | ~5.5 Å → ~13.8 Å<br>($\Delta \approx 8.3$ Å) | ~7.2 Å → ~16.2 Å<br>($\Delta \approx 9$ Å) |
| N525–K616 | 4.36–6.38 | ~ 11.0 Å → ~ 18.0 Å<br>($\Delta \approx 7.0$ Å) | ~12.2 Å → ~ 18.8 Å<br>($\Delta \approx 6.6$ Å) | ~ 13.4 Å → ~ 20.5 Å<br>($\Delta \approx 7.1$ Å) |

This comparison reveals that TMEM16F undergoes a larger opening at the extracellular entrance of its groove than the fungal homologs, as reflected in the substantially greater increase in the 4.21–6.26 pair distance. The GNS thus enables a quantitative, position-by-position comparison of groove-opening geometry across phylogenetically diverse TMEM16 scramblases.

##### Mutagenesis of Groove-Lining Residues

A further illustration of the GNS is provided by a comparison of mutagenesis data for nhTMEM16 and afTMEM16. Using the identification of corresponding positions, we found that individual mutations of groove-lining residues shown to impair substantially the scramblase function in nhTMEM16, appear to have little or no effect at the equivalent positions in afTMEM16 (Falzone et al., 2022; Lee et al., 2018). For example, the E313–R432 salt bridge in nhTMEM16 has been shown by mutagenesis to be important for scrambling activity, whereas the equivalent E305–R425 pair in afTMEM16 does not contribute comparably to function.

We also observed a similar difference in the response to mutations by the two fungal scramblases in the results of Alanine substitutions in residue pair *E3.55(313)-R6.26(432)* in nhTMEM16 compared to such substitutions in the corresponding afTMEM16 residues *E3.55(305)* and *R6.26(425)*. Despite the equivalent structural locations, the same experimental laboratory (Falzone et al., 2022; Lee et al., 2018) found that Ala substitution of the nhTMEM16 E–R pair caused a greater than 100-fold reduction in scramblase activity, whereas the corresponding substitution in afTMEM16 produced less than a 2-fold change. This result highlights an important limitation of any GNS: structural equivalence of position does not guarantee functional equivalence of the residue. The GNS is a tool for identifying corresponding loci; whether those loci perform equivalent functions must be established experimentally.

##### Considerations for Applying the Numbering Scheme to New Sequences

Given the large number of TMEM16 subtypes and the relatively limited number that have been experimentally characterized, we considered the need for a way to extend the GNS to sequences not represented in the reference alignment. Were a new sequence with low similarity to be added to the TMEM16-RA by sequence alignment alone, it may lead to incorrect TsRR assignments (e.g.,by introducing spurious gaps within TMs). We propose to address this by combining structural alignment with the sequence alignment.

Specifically, for TMEM16 subtypes with experimentally determined structures, the new structure can be aligned first to mTMEM16F for which structure-function information is available. If the resulting sequence alignment of the reference homologs and the new subtype place the TsRR residues at the same alignment positions as in the existing TMEM16-RA, that alignment is accepted and the numbering is assigned directly from the reference positions (N.50). If the TsRR positions shift in the new alignment, structural superposition is used to determine which residues occupy the same spatial positions as the TsRRs in mTMEM16F; further analysis relative to functional information can then be used before the numbering scheme can be applied with confidence.

For TMEM16 subtypes lacking experimental structural information, the same procedure can be applied using predicted structures generated by AlphaFold (Jumper et al., 2021). Although N- and C-terminal regions are often poorly modeled by AlphaFold, the transmembrane helices are generally well predicted, enabling reliable identification of residues occupying positions equivalent to the TsRRs in mTMEM16F. Note that all TMEM16 structures in the AlphaFold Protein Structure Database (Varadi et al., 2022) correspond to the closed-groove conformation; accordingly, the closed-groove conformation of mTMEM16F is the recommended reference for structural alignment in this protocol.

## Conclusions

We have developed a reference alignment (RA)-based generic numbering scheme (GNS-TMEM16) for the TMEM16 family of phospholipid scramblases, providing a unified positional reference framework analogous to the long-established Ballesteros and Weinstein numbering system for class A GPCRs (Ballesteros & Weinstein, 1995). The scheme is built on a curated reference alignment (TMEM16-RA) of twelve human and mouse TMEM16 scramblases—TMEM16C, D, E, F, G, and J—chosen for their high degree of sequence identity and the availability of experimentally determined structures. As described, this reference alignment is further augmented by a small number of sequences that are phylogenetically somewhat more distant, resulting in an augmented reference alignment (TMEM16-ARA) that contains a group composed of the mammalian TMEM16K, the fungal scramblases afTMEM16 and nhTMEM16, and the TMEM16 channels TMEM16A, and TMEM16B. This TMEM16-ARA is used to identify a TM-specific reference residue (TsRR) for each of the ten transmembrane helices through hierarchical application of three criteria: 100% conservation within the core TMEM16-RA (Criterion 1); additional conservation within at least one of the analogs in the augmenting group (Criterion 2); and structural and functional considerations including helix-perturbing character, groove localization, conserved motif membership, and central position within the TM (Criterion 3). The ten TsRRs identified by this protocol span a range of physicochemical properties— Tyr (TM1), Trp (TM2), Arg (TM3), Glu (TM4), Phe (TM5), Pro (TM6), Glu (TM7), Asp (TM8), Trp (TM9), and Glu (TM10). These TsRRs include residues that are specifically implicated in Ca²⁺ coordination (TMs 6–8), groove lining, and helix structural stabilization.

Residues in each TM are assigned an identifier of the form **N.m(k)**, where **N** is the TM number, **m** is the residue’s position relative to the TsRR (which assigned m = 50), and **k** is the absolute sequence number in the specific protein. Residues in TM-connecting loops receive dual identifiers measured from the TsRRs of both flanking helices (e.g., D3.70/4.08(497)), and mutations are represented in the standard format (e.g., R6.26(432)W). Residues in the N- and C-terminal regions outside the TM bundle are excluded from the scheme.

By using the consistent numbering scheme across subtypes, we illustrate the use of the GNS-TMEM16 and demonstrate its immediate utility in identifying various conserved structural components of the scramblases that support their functional mechanisms. The first demonstrates a systematic, position-by-position comparison of groove-opening geometry across phylogenetically diverse scramblases.

Translating the four TM4–TM6 residue pair distances characterized by Khelashvili et al. (Khelashvili et al., 2022) in mTMEM16F into equivalent positions in afTMEM16 and nhTMEM16, reveals that the extracellular entrance of the groove (pairs 4.21–6.26 and 4.25–6.30) opens substantially more in mTMEM16F than in either fungal homolog (consistent with the distinct helix-loop-helix cap architecture of its TM2), and is wider in nhTMEM16 than in the afTMEM16 homolog. This comparison would be impractical without a shared positional reference, since the absolute sequence numbers of equivalent residues differ across homologs. In the second application, the GNS identifies structurally equivalent positions across paralogs in the context of functional divergence. The E3.55–R6.26 salt bridge, whose disruption by Ala substitution reduces scramblase activity more than 100-fold in nhTMEM16 (Falzone et al., 2022; Lee et al., 2018), occupies the equivalent locus in afTMEM16 (E3.55(305)–R6.26(425)), yet the same mutations produce less than a 2-fold functional change in that homolog. This result illustrates a critical interpretive point: the GNS identifies structural correspondence of position, but not functional equivalence of the residue at that position—a distinction that must be resolved by experiment and that the GNS is specifically designed to expose and facilitate.

We also provide a protocol for extending the GNS-TMEM16 to sequences not represented in the reference alignment, combining ClustalW sequence alignment with structural superposition to the mTMEM16F closed-groove conformation (PDB: 6QPB). For sequences lacking experimental structures, AlphaFold-predicted models (Jumper et al., 2021) can be substituted, as TM helices are reliably modeled even when terminal regions are not. This protocol makes the GNS-TMEM16 immediately applicable to the full breadth of characterized and uncharacterized TMEM16 family members.

More broadly, inspection of the TMEM16-RA reveals that the most highly conserved TMs (i.e., 6, 7, and 8) are those that coordinate Ca²⁺ binding, while TM3, which lines the protomer groove and directly contacts phospholipid substrates during scrambling, is among the least conserved. This pattern suggests that the Ca²⁺-sensing apparatus is under stronger evolutionary constraint than the groove-lining surface, implying that functional diversification across the family may be achieved in part through variation in groove composition while the activation mechanism is conserved. The GNS-TMEM16 provides the positional framework needed to test this hypothesis systematically as structural and functional data continue to accumulate for the broader TMEM16 family.

Finally, the three-criterion selection procedure developed here anchors a GNS to 100%-conserved residues identified from a curated reference alignment that is refined by phylogenetically extended conservation and structural considerations. It is general and can be applied to other polytopic membrane protein families that share a common transmembrane architecture. For any such family, the relatively rigid transmembrane helices provide the structural pivots needed to establish a universal positional reference, enabling the kind of cross-subtype comparisons that have proven transformative in GPCR biology.

## Methods

### Sequence Acquisition

All TMEM16 sequences were retrieved from UniProtKB (Coudert et al., 2023; UniProt Consortium, 2023): hTMEM16A (Q5XXA6), mTMEM16A (Q8BHY3), hTMEM16B (Q9NQ90), mTMEM16B (Q8CFW1), hTMEM16C (Q9BYT9), mTMEM16C (A2AHL1), hTMEM16D (Q32M45), mTMEM16D (Q8C5H1), hTMEM16E (Q75V66), mTMEM16E (Q75UR0), hTMEM16F (Q4KMQ2), mTMEM16F (Q6P9J9), hTMEM16G (Q6IWH7), mTMEM16G (Q14AT5), hTMEM16J (A1A5B4), mTMEM16J (P86044), hTMEM16K (Q9NW15), mTMEM16K (Q8BH79), afTMEM16 (Q4WA18), nhTMEM16 (C7Z7K1).

### Structure Acquisition

All structures were downloaded from the Protein Data Bank (Berman et al., 2000); PDB accession codes are cited in the main text.

### Sequence Alignment and Phylogenetic Analysis

Multiple sequence alignment and phylogenetic tree construction were performed in MEGA11 (Tamura et al., 2021), following the protocol described by Hall (Hall, 2013) (Steps 2 and 3). Alignment was performed using ClustalW (Thompson et al., 1994), which produced fewer gaps and more consistent conservation patterns across the selected TMEM16 sequences than MUSCLE, and whose outputs were most consistent with existing functional analyses. The Multiple Alignment Gap Opening Penalty was set to 3 and the Gap Extension Penalty to 1.8. The best-fit substitution model was selected automatically using the Find Best DNA/Protein Models (ML) function in MEGA11; the LG + G model was chosen.

All subsequent alignments for TsRR selection were performed using Clustal Omega with default settings (Sievers et al., 2011; Sievers & Higgins, 2018).

## Acknowledgments

The authors thank Drs. Alessio Accardi and George Khelashvili for helpful discussions and gratefully acknowledge the support of this research by a grant from the 1923 Foundation (to HW). Computational resources of the David A. Cofrin Center for Biomedical Information in the HRH Prince Alwaleed Bin Talal Bin Abdulaziz Alsaud Institute for Computational Biomedicine at Weill Cornell Medical College, and the resources under Project BIP109 at the Oak Ridge Leadership Computing Facility, which is a DOE Office of Science User Facility supported under Contract DE-AC05-00OR22725 are gratefully acknowledged.

## Supplementary Information

**Supplement Figure 1.**
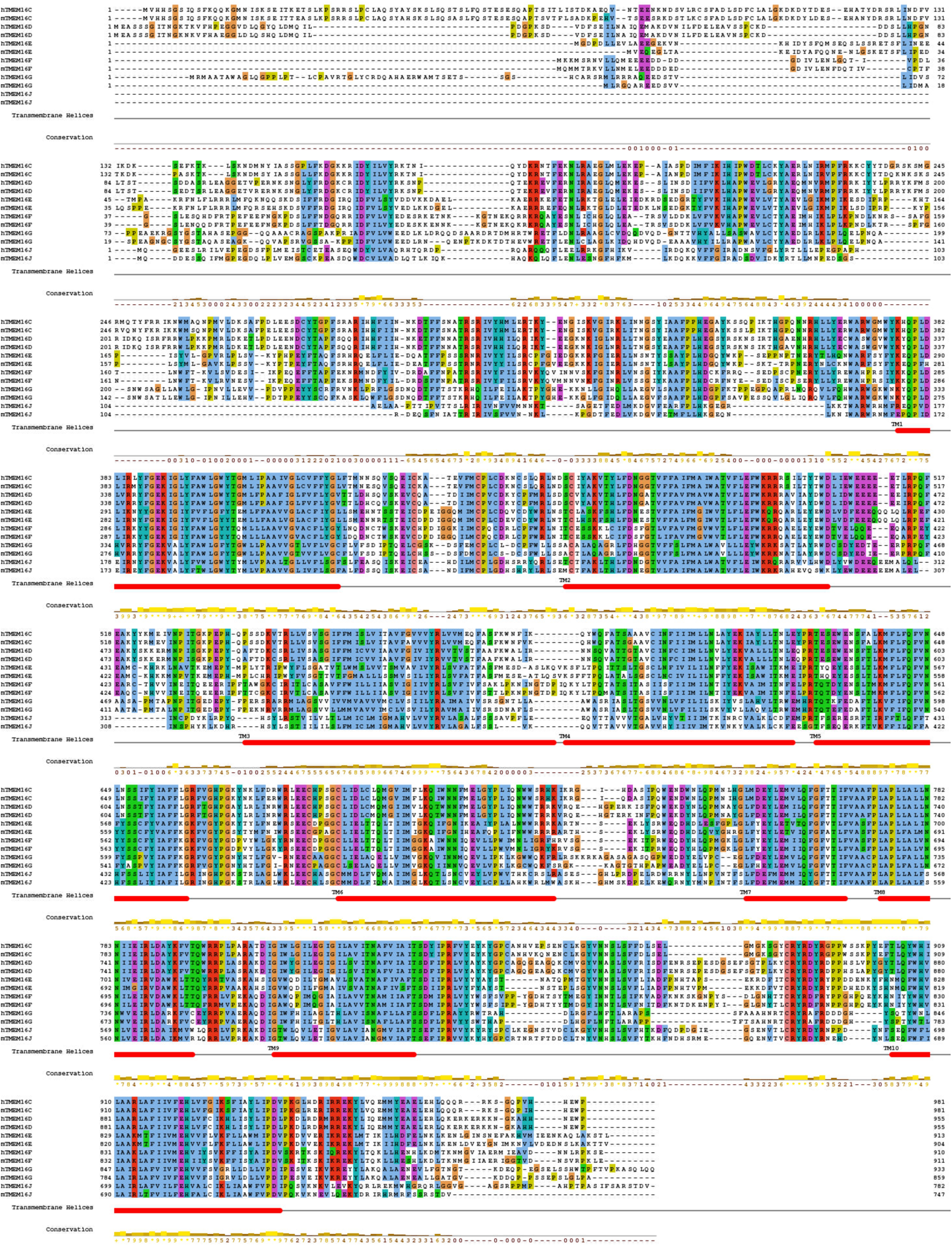
Reference alignments (TMEM16-RA) of the TMEM16 sequences selected for the reference set as described in the text. The alignment includes the sequences of human and mouse TMEM16 proteins C, D, E, F, G, and J which are considered to function as phospholipid scramblases and is illustrated with Jalview (Waterhouse et al., 2009). The TM regions defined according to the specified segments in these PDB structures, are indicated by the thick horizontal red underlines. The color scheme for the residues is defined by ClustalX. The conservation percentage is calculated automatically by Jalview, and 100% conservation is identified by the vertical red lines in the helical segments.

